# Transient DNA binding to gapped DNA substrates links DNA sequence to the single-molecule kinetics of protein-DNA interactions

**DOI:** 10.1101/2022.02.27.482175

**Authors:** Rebecca Andrews, Horst Steuer, Afaf H. El-Sagheer, Abhishek Mazumder, Hafez el Sayyed, Arun Shivalingam, Tom Brown, Achillefs N. Kapanidis

## Abstract

Protein interactions with nucleic acids are central to all genetic processes and many biotechnological applications. While many sequence-dependent protein-DNA interactions have been studied in detail using single-molecule methods, there is no standard high-throughput way to link the complex single-molecule kinetics of protein-DNA interactions with the DNA sequence of a single molecule. Here we provide the missing link by introducing a single-molecule imaging method (Gap-Seq) that interrogates DNA sequences via transient binding of short fluorescent DNA to a single DNA molecule previously used to characterise a protein-DNA interaction. In Gap-Seq, we identify a base by the degree of binding of 6-9 nt-long DNAs to surface-immobilised DNA substrates featuring a short single-stranded gap. To facilitate detection, we also developed a fluorescence quenching strategy that allows single-molecule detection at up to 500 nM of unbound fluorescent DNA. We link single-base differences on single DNA molecules to the kinetics of protein-DNA interactions by studying the interaction of a transcription activator with its cognate site. Finally, we show that our assay can address mixed sequences by distinguishing between two different sequences immobilised on the same field of view, paving the way for interrogation of sequence libraries for both mechanistic work and biotechnological applications.

## INTRODUCTION

Sequence-specific interactions of biomolecules, especially proteins with DNA and RNA, lie at the heart of all processes in the storage, maintenance and expression of genetic information. The analysis of such interactions relies on structural approaches that provide detailed views, often at the atomic level, of the complexes involved. In addition, biochemical and biophysical approaches report on thermodynamic and kinetic aspects of these interactions. In particular, the wide availability of second-generation sequencing over the past decade offered a systematic, high-throughput and high-resolution avenue for the analysis of protein-DNA and protein-RNA interactions both *in vitro* and *in vivo* (1–3).

However, the use of genome sequencing approaches to dissect protein interactions with nucleic acids is complicated by ensemble averaging. Such approaches cannot report directly on the kinetics of protein-DNA interactions, especially when these interactions involve several steps that go beyond relatively simple binding-unbinding equilibria, as in the case of multi-step reactions in gene repair, editing, replication, and transcription, all of which feature transient on-pathway or off-pathway intermediates (4–9). NGS methods also typically rely on DNA amplification, introducing the possibility of base errors.

On the other hand, single-molecule analysis of sequence-specific protein-DNA and protein-RNA interactions can uniquely address issues of sample heterogeneity, while offering the ability to monitor interactions and associated conformational changes in real-time (4–6, 10, 11). Such methods include single-molecule fluorescence, force-based methods, and nanopore-based methods (10). However, there is currently no standard high-throughput single-molecule method that can assess the sequence-specificity of protein-DNA interactions; typically, single-molecule measurements interrogate one sequence at a time, and assess sequence dependence by the comparison of the behaviour on only a few sequences assessed in separate experiments. As a result, it is difficult to harness the power of single-molecule methods towards a systematic analysis of the protein-DNA interactions, their structural features, and their kinetics.

A possible avenue towards high-throughput analysis of the sequence-dependence of protein-DNA interactions involves the coupling of existing commercial single-molecule sequencing methods (nanopore-based, or single-molecule-fluorescence-based) to kinetic analysis of protein-driven processes on the strands of the DNA that are being sequenced (12–14). Within the last decade, single-molecule sequencing has become established in the commercial long read sequencing market through the offerings of companies such as Pacific Biosciences and Oxford Nanopore Technologies. However, these approaches either require expensive specialized equipment and special surfaces (e.g., zero-mode waveguides in the case of single-molecule fluorescence (12)) or are largely confined to the machines that translocate on nucleic acids, such as DNA helicases and RNA polymerases (14). As a result, these methods do not offer a general approach to deciphering the sequence-dependence of most protein-DNA interactions.

Here, we introduce a new approach towards the systematic analysis of protein-DNA interactions. Our approach relies on connecting the functional properties of a single DNA (or RNA) molecule (its “single-molecule phenotype”) with its DNA sequence; the functional properties may involve structure, dynamics, recognition/binding by a protein, and processing by a protein. Briefly, surface-immobilised DNAs containing sequence segments that can be randomized (to provide libraries that represent a large pool of sequences) are interrogated with regards to their interaction with a protein using single-molecule fluorescence imaging, and then the sequence of each DNA molecule is connected to the observed single-molecule phenotype.

To link the single-molecule phenotype to DNA sequence, we introduce a method that reports on the sequence of single DNA molecules based on transient hybridization of short DNA strands (6-9 nt) to surface-immobilised DNA; such transient binding has also been exploited in other high-throughput endeavours, such as super-resolution imaging by DNA-PAINT (15). The sequence identification relies on the fact that the binding of a short but fully complementary DNA is more stable than that of a DNA having the same length but including one or more mismatches to the surface-immobilised DNA. Since our approach involves DNAs with single-stranded gaps, we characterize the binding of short DNA to these substrates, and show that we can detect single nucleotide differences on a DNA template by the binding of short DNA strands ranging from 6 to 9 nucleotides in length. Finally, we link the sequence of a single DNA molecule with its interaction with a sequence-specific DNA binding protein, and provide proof-of-concept for the identification of sequence variants within in a mixed sample of multiple DNA sequences immobilised randomly on the surface. Our results also inform on the mechanisms and kinetics of hybridization and have implications for DNA nanotechnology.

## METHODS

### Preparation of DNA fragments

Oligodeoxyribonucleotides (“oligos”) were predominantly prepared in house (see below for detailed description), with the remaining oligos coming from commercial sources (IBA Life Sciences; IDT); in the latter case, the oligos were dissolved at a final concentration of 100 μM and stored at −20°C. The full list of all oligos used is in Table 1.

**Table 1:**
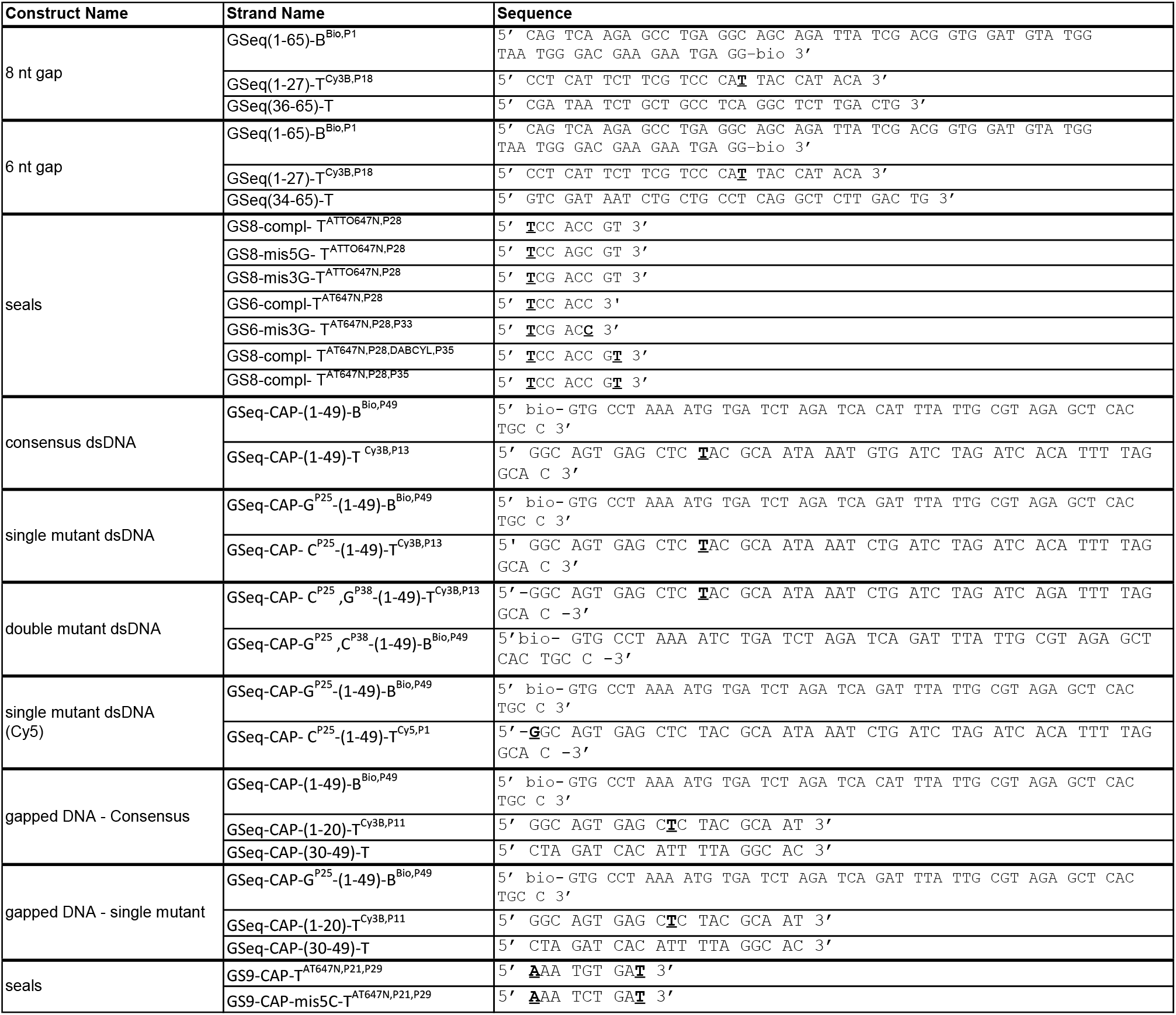
DNA sequences for all constructs and seals. In bold typeface: locations of fluorophores and quenchers.

Unmodified and modified oligonucleotides prepared in house were synthesized on an Applied Biosystems 394 automated DNA/RNA synthesizer using a standard 1.0 μmole phosphoramidite cycle of acid-catalyzed detritylation, coupling, capping, and iodine oxidation. Stepwise coupling efficiencies and overall yields were determined by the automated trityl cation conductivity monitoring facility and was >98.0%. Pre-packed SynBase™ CPG 1000/110 (Link Technologies) 3′-amino C7, 3′-biotin, 3′-dabcyl and 3′-Cy3 resins were used for the synthesis. All β-cyanoethyl phosphoramidite monomers were dissolved in anhydrous acetonitrile to a concentration of 0.1 M immediately prior to use with coupling time of 50 s for normal A, G, C, and T monomers and was extended to 600 s for 5’-amino, 5’-Cy3, 5’-biotin and 5-amino C6 dT phosphoramidite monomers (from Link Technologies Ltd.).

5’-Amino, 3’-amino and internal 5-amino C6 dT modified oligonucleotides on resin were treated with a solution of diethylamine (10% in acetonitrile) for 20 min in order to selectively remove the cyanoethyl protecting groups. Cleavage and deprotection of oligonucleotides were achieved by exposure to concentrated aqueous ammonia solution for 60 min at room temperature followed by heating in a sealed tube for 5 hrs at 55°C.

Oligonucleotides labelled with either: 1) two ATTO647N fluorophores; 2) an ATTO647N fluorophore and a DABCYL quencher; or 3) Cy3B were prepared by NHS-ester coupling. The unlabelled pre-cursor oligonucleotide (40 nmol) was freeze-dried and resuspended in sodium carbonate buffer (18 μl, 0.5 M, pH 8.5). Separately, Cy3B or ATTO647N NHS-ester (6 μl, 20 nmol/μl in DMSO) was mixed with DMSO (6 μL). The NHS ester solution was then added to the oligonucleotide solution and mixed vigorously before shaking (800 rpm, room temperature). After 3 hrs, reactions were diluted with water (970 μl) and purified by desalting on a disposable gel-filtration column (NAP10, GE Healthcare Life Sciences, cat. no. GE17-0854-02) according to the manufacturer’s instruction to remove the majority of the free dye. Labelled oligonucleotides were further purified by reverse-phase HPLC chromatography using a Gilson HPLC system with ACE® C8 column (10 mm x 250 mm, pore size 100 Å, particle size 10 μm) with a gradient of buffer A (0.1 M TEAB, pH 7.5, where TEAB = triethylammonium bicarbonate) to buffer B (0.1 M TEAB, pH 7.5 containing 50% v/v acetonitrile) and flow rate of 4 ml/min over a total run time of 17.5 min. For all ATTO647N labelled oligonucleotides, a gradient of 70-100%buffer B was used. For Cy3B-labelled oligonucleotides, a gradient of 20-35% buffer B was used. For other oligonucleotides, a gradient of 10-25% buffer B was used. The TEAB buffer and acetonitrile in the collected fractions were removed *in vacuo* or by freeze drying.

All oligonucleotides were characterised by negative-mode electrospray using a UPLC-MS Waters XEVO G2-QTOF mass spectrometer and an Acquity UPLC system with a BEH C18 1.7 μm column (Waters). A gradient of methanol in triethylamine (TEA) and hexafluoroisopropanol (HFIP) was used (buffer A, 8.6 mM TEA, 200 mM HFIP in 5% methanol/water (v/v); buffer B, 20% v/v buffer A in methanol). Buffer B was increased from 0–70% over 7.5 min or 15–30% over 12.5 min for normal oligonucleotides and 50–100% over 7.5 min for hydrophobic oligonucleotides. The flow rate was set to 0.2 mL/min. Raw data were processed and deconvoluted using the deconvolution software MassLynx v4.1 and in all cases confirmed the integrity of the sequences.

With regards to our DNA nomenclature, in a strand named GSeq(1-27)-T^Cy3B,P18^, GSeq refers to gap sequencing experiments, (1–27) refers to the length of the strand and the position within the construct with respect to the 5′ end, T denotes the top strand, and the superscript (Cy3B, P18) notes the DNA modification and its position.

To form the gap-containing DNA substrates, as well as the fully duplexed DNA fragments, oligos were annealed by mixing two complementary strands in a ratio of 1:1 in hybridization buffer (200 mM Tris–HCl pH 8.0, 500 mM NaCl, 1 mM EDTA) and by heating for 5 min at 95°C, followed by cooling to 25°C at a rate of 2°C per minute and placed on ice before being used.

### Preparation of labelled CAP derivative (Alexa647-CAP)

Overexpression and purification of the catabolite activator protein CAP-(Cys17, Ser178) was performed using plasmid pHSCRP-His6-H17C-C178S [constructed from plasmid pAKCRP-His6 (16) using site-directed mutagenesis (Q5 Site-Directed Mutagenesis Kit, NEB)].

Fluorescence labelling was performed by incubating 20 μM CAP-(Cys17, Ser178) with 400 μM Alexa647 maleimide in buffer A [40 mM HEPES-NaOH (pH 7.3), 100 mM NaCl, 2 mM TCEP and 5% glycerol] for 12 hrs on ice. The fluorescent probe labelled protein (Alexa647-CAP) was purified using a G25 Sephadex gel filtration column pre-equilibrated with buffer B (40 mM Tris-HCl pH 8, 100 mM NaCl, 0.1 mM TCEP and 5% glycerol), followed by buffer exchange (5 cycles of concentration to 0.5 ml volume and dilution with 5 ml buffer B) using a 3 kDa MWCO Amicon Ultra-15 centrifugal ultrafilter (Millipore), and the purified Alexa647-CAP was stored in aliquots at −80°C.

### Sample preparation for real-time transient DNA binding

The biotinylated DNA constructs were immobilised at ~0.2 μm^-2^ density via neutravidin, inside the wells of silicon gaskets, on plasma-cleaned coverslips coated with polyethylene-glycol (PEG; (4, 17)). We then added 20 μl of 50 pM gapped-DNA constructs in Milli Q, incubated for 10 s, and washed three times with 200 μl PBS. Subsequently, we added 20 μl of DNA imaging buffer (0.5 M NaCl, 50 mM HEPES pH 7.3, 6 mM BSA, 1 mM TROLOX, 1% Glucose, 40 μg/ml catalase and 0.1 mg/ml glucose oxidase) containing 10 nM (unquenched) or 100 nM (quenched) DNA seals in the well.

### Sample preparation for real-time CAP-DNA interactions

DNA fragments containing derivatives of the consensus sequence for CAP binding were prepared as above. CAP was added at 1.5 nM CAP concentration and binding was imaged in the presence of the CAP binding buffer (0.2 mM cAMP, 40 mM Tris pH 8, 100 mM KCl, 10 mM MgCl_2_, 5% glycerol, 40% glucose and 1 mM TROLOX) plus an oxygen-scavenging system (1 % Glucose, 40 μg ml^-1^ catalase and 0.1 mg ml^-1^ glucose oxidase).

### In situ preparation of gapped-DNA substrates

After monitoring the CAP-DNA interactions on immobilised DNA molecules via single-molecule imaging, the sample was washed three times with 200 μl 1xPBS, and then incubated with 25 mM of NaOH for one minute. After washing again with 200 μl 1xPBS, imaging was performed in DNA imaging buffer to ensure the completion of DNA denaturation. The sample was then washed with 200 μl 1xPBS, followed by 5 min incubation with 1 nM GSeq-CAP-(1-20)-T^Cy3B,P11^ and 1 nM GSeq-CAP-(30-49)-T in annealing buffer. The sample was then washed three times with 200 μl 1xPBS and imaged in the presence of DNA imaging buffer and seal DNA at 100 nM.

### Mixed DNA on the surface

In Step 2, the NaOH wash was performed for 2 s instead of 2 min and the gap hybridisation mix contained 100 nM of the top flanking stands. We also included surface-immobilised 60-nm gold colloid nanoparticles (EM.GC60, BBI Solutions; diluted 1:100 in MilliQ water and sonicated for 10 minutes before being placed on surface) to act as fiducial markers in data analysis for correction of the lateral drift of the field of view.

### Single-molecule imaging

Single-molecule fluorescence movies were collected using the Nanoimager-S single-molecule fluorescence microscope (Oxford Nanoimaging; (18, 19)). The microscope was used as a widefield single-molecule fluorescence microscope with objective-based total internal reflection fluorescence (TIRF) illumination mode, with an excitation angle set at 52°. We performed our imaging using continuous-wave excitation (532 nm for Cy3B and 640 nm for ATTO647N and Alexa647), with typical laser powers of ~0.4 mW for 532-nm excitation and ~45 mW for 640-nm excitation. In all experiments, we used 100-ms exposures, apart from the detection of the gap binding of 6-nt seals, where 10-ms exposures were used; typical traces were 5 min long, with occasional movies being 10 min long. The measurements were performed at microscope temperatures of 27-30°C, as indicated by the Nanoimager temperature sensor. In experiments where a biotinylated gapped-DNA was labelled by Cy3B, we initially used 532-nm laser excitation to localise of the gapped DNA, and then switched to 640-nm excitation to directly excite the ATTO647N-labelled seal DNA upon binding to the gapped-DNA.

### Time-trace analysis and dwell-time distributions

The raw image was processed by home-built software (GapViewer) that detects diffraction-limited spots by their intensity (18). The software performs local background subtraction for each frame using the intensity around the Gaussian profile of the fluorescence emission from single DNA molecules. The locations of the spots were determined from a maximum-intensity projection of the raw movies by finding local maxima in the background subtracted signal. The resulting time-trace data in the red and green channels were sorted manually to ensure the traces correspond to single localisations. Following Hidden Markov Modeling (HMM) analysis of the traces, and the removal of events lasting less than three frames, the dwell-times for the DNA-bound and unbound states were extracted, and plotted as frequency histograms. For the DNA-bound states, we used a bin size of 10 frames (i.e., 1-s bins for movies with 100-ms exposures, and 100-ms bins for movies with 10-ms exposures); for the off-time histograms (corresponding to the unbound states), we used a bin size of 5 s. We fitted our dwell-time distributions in Origin (OriginLab) using either a single- or double-exponential decay. During the analysis of the mixed-DNA samples, correction of lateral drift using fiducial markers was done using the “Localize” and “Render” functions of the Picasso software (20).

## RESULTS

### Concept of linking single-molecule phenotypes to DNA sequence

Our motivation for the work was the development of a high-throughput assay (Fig. 1) that links single-molecule phenotypes (as they apply to sequence-dependent protein-DNA interactions) to DNA sequence for libraries of DNA fragments.

**Figure 1:**
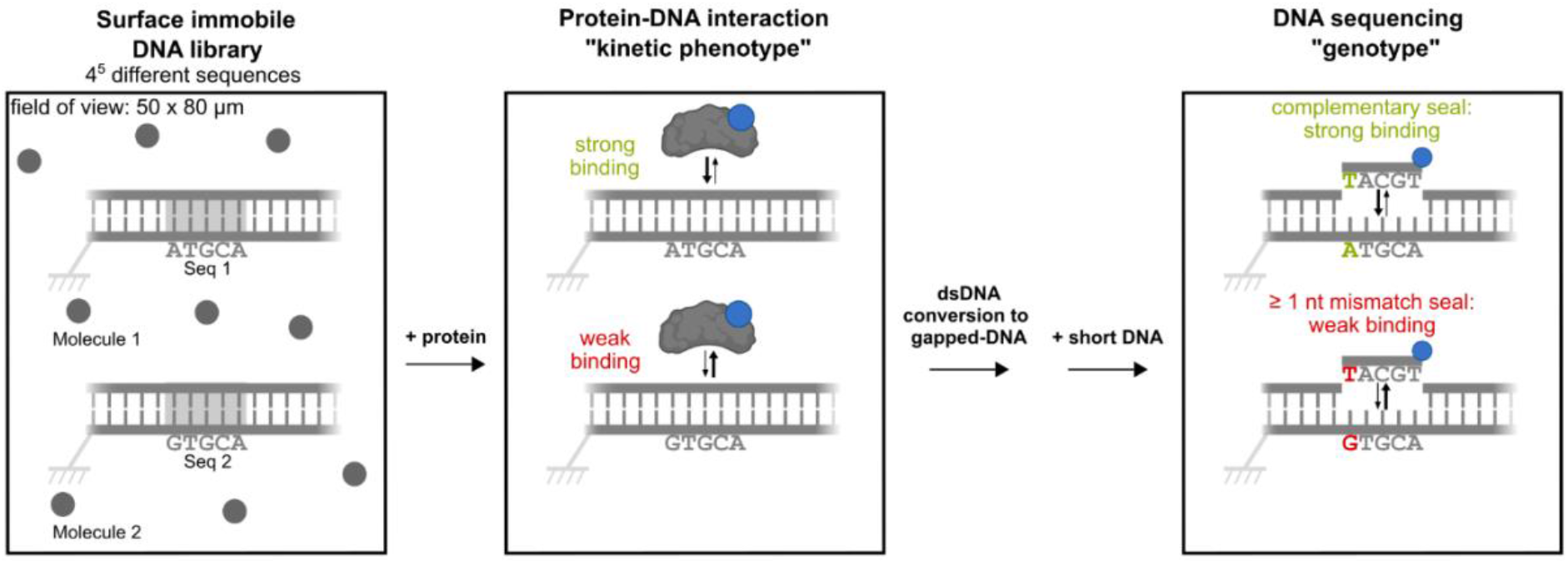
A single-molecule method that links protein-DNA interactions to DNA sequence. Left panel: example of a field-of-view showing 9 diffraction limited images (gray circles) of single DNA molecules immobilised on a glass surface and imaged using a widefield fluorescence microscope. The DNA molecules form part of a library with 4^5^ different DNA sequences. Middle panel: fluorescently labelled proteins of interest are added, and the single-molecule kinetics of protein-DNA interactions (its “single-molecule kinetic phenotype”) are recorded. Transition to right panel: double-stranded DNA is denatured, leaving the immobilised single strand on the surface; gapped DNA molecules are assembled *in situ* on the ssDNA on the surface, and short fluorescent oligonucleotides (seals) are added. Right panel: DNA seals bind to the gap in the surface-immobilised gapped-DNA, with the kinetics of binding discriminating between fully complementary seals and those with one or more mismatches.

The first step of the assay involves generating a library of DNA molecules representing different sequences, and immobilising these DNAs randomly on a glass surface (Fig. 1, left; the rectangle represents an example field-of-view). In the second step, a fluorescently labelled protein of interest is introduced, and the protein-DNA interactions are examined (Fig. 1, middle), providing a single-molecule phenotype (also referred to as a “single-molecule kinetic phenotype”, when kinetic information is extracted) which can be much more complex than simple binding-unbinding equilibria. In the third step (Fig. 1, right), the protein is removed, the immobilised DNA is prepared for sequence interrogation, and a single-molecule fluorescence readout reports on the DNA sequence of each single molecule.

Steps 1 and 2 in our scheme are well established within single-molecule biophysics; however, no standard approach exists for step 3. We thus introduce a method capable of examining short DNA sequences using transient binding of fluorescent single-stranded DNA oligos (“seals”) to surface-immobilised DNAs that feature short single-stranded DNA gaps (Fig. 2A); we hence refer to this method as “Gap-Seq”.

**Figure 2:**
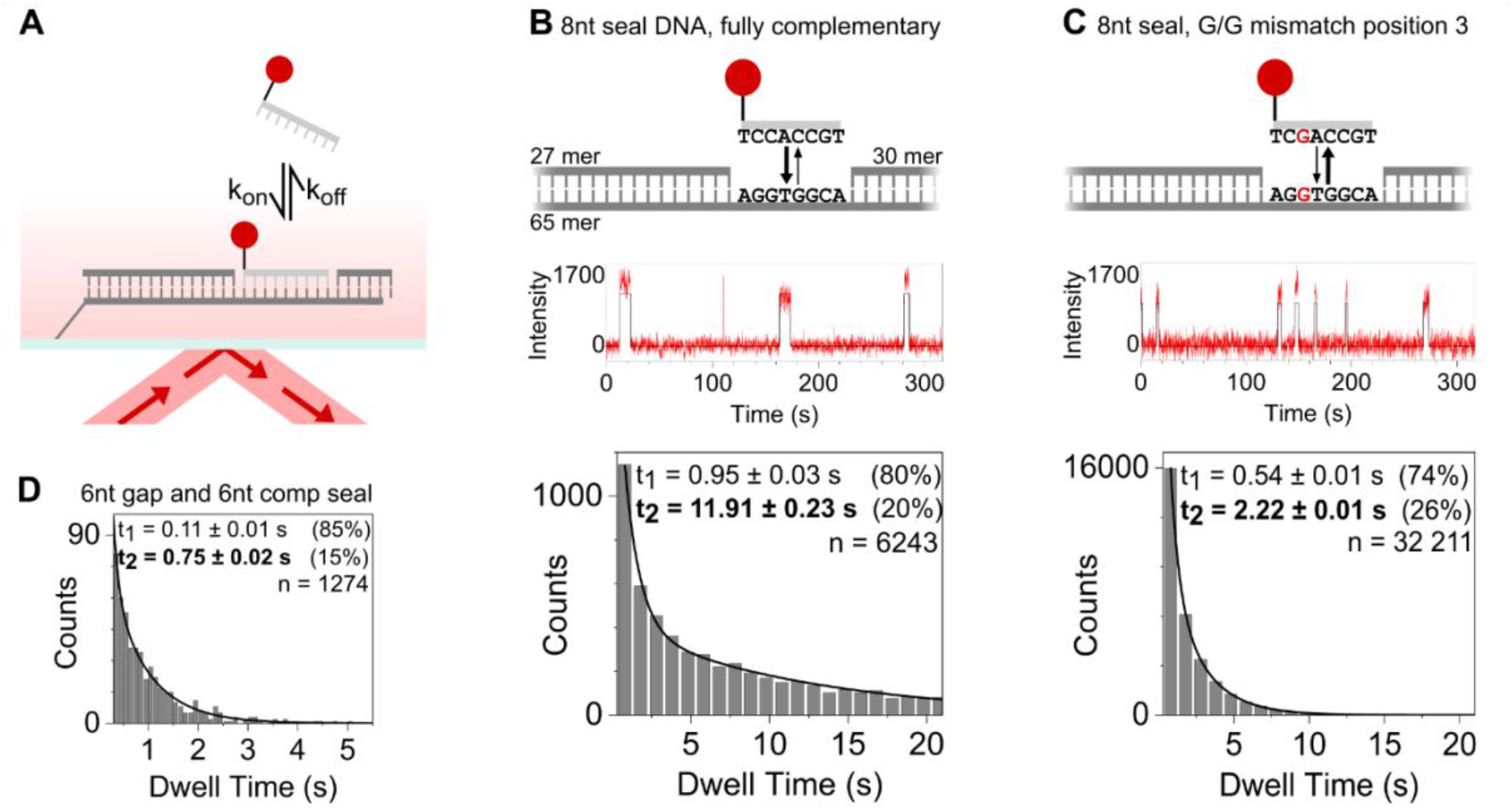
Detecting single base-pair mismatches via transient DNA binding to gapped DNA. A. Schematic showing the transient binding of labelled oligonucleotides (seals) to surface-immobilised gapped DNA; detection is performed via total internal reflection fluorescence (TIRF) microscopy. B. Top: schematic of a fully complementary 8-nucleotide (nt) seal binding in 8-nt gapped-DNA. Middle: fluorescence intensity time-traces (red) for ATTO647N-labelled DNA seal binding to gapped DNA. Movies recorded using 100-ms exposures. Traces are fitted (black) using Hidden Markov Modelling (HMM), identifying bound states and measuring the corresponding dwell times. Bottom: dwell-time distribution for the bound states. The data fit best to double-exponential distributions that correspond to a long and short dwell time. n: number of binding events. C. As in panel B, but for a G-G mismatch at position 3 in the 8-nt seal binding to an 8-nt gapped-DNA. D. As in panel B, but for fully complementary 6-nt seal binding in a 6-nt gapped-DNA. Movies recorded using 10-ms exposures. In contrast, no binding events were detected for the same observation time using a 6-nt seal with a mismatch at position 3.

The short length of the gap (6-9 nt) allows the bound seals to thermally dissociate and quickly be replaced by other DNA seal molecules from the surrounding solution; repeated binding of the seals provides increased statistics that allows base interrogation with higher accuracy, while eliminating photobleaching through the abundant supply of unbleached seals. Such transient binding is also used in the super-resolution imaging technique DNA PAINT (15), which uses ssDNA as a binding template. In our work, we selected to use gapped-DNAs instead of ssDNA, since the coaxial stacking provided by the bases at the end of the gap (not present in ssDNA) helps to stabilise the bound seal; further, the presence of a gap defines more clearly the short DNA segment to be interrogated. Analysis of the extent of repeated seal binding should allow us to distinguish between DNA seals which are fully complementary, and DNA seals with one or more base-pair mismatches.

### Reading single base information from transient DNA binding to an 8-nt gapped DNA

To establish our ability to detect single nucleotide differences within immobilised gapped DNA, we first tested our ability to detect a single base mismatch between the gapped-DNA substrate and transiently binding seals.

Our first experiments were performed using surface-immobilised gapped-DNA with an 8-nt gap, and 8-nt-long seals fully complementary to the gap (Fig. 2A). After localising the gapped-DNAs using Cy3B fluorescence (placed on the top strand flanking the gap on left side), we detected the seal DNA binding via direct excitation of a red fluorophore (ATTO647N, placed at the 5’-end of the seal DNA) during the short dwell of the seal DNA on the immobilised strand (Fig. 2A). To monitor multiple binding events of the seals while keeping background fluorescence low, seals were kept at 10 nM in the solution surrounding the gapped-DNAs. Upon seal binding, we observe a spike in the fluorescence intensity which lasts until the seal dissociates, at which point the fluorescence intensity returns to the level of background (Fig. 2B); the fluorescence intensity time-trace in Fig. 2B shows four high fluorescence intensity events (lasting from ~100 ms to 15 s), which correspond to seal binding to an individual gapped-DNA molecule.

After measuring the dwell times for the repeated seal binding to each gapped-DNA molecule in the field of view (FOV), we plotted the dwell-time distribution for the binding of an 8-nt fully complementary seals to gapped DNA (*n* = 6,243; Fig. 2B, bottom). The distribution was fit poorly by a single-exponential decay distribution (Fig. S1), but was described very well by a double-exponential decay with dwell times t_1_ and t_2_. We reasoned that the shorter dwell time, t_1_ ~ 0.95 s (accounting for ~80% of the events) corresponds to partial hybridisation and rapid dissociation of the seal DNA (see later sections and *Discussion*), whereas the longer time, t_2_ ~ 11.9 s (accounting for ~20% of the events), corresponds to the binding time of a fully bound, fully stacked seal; since the latter is a more accurate reporter for complete binding (both due to the length and nature of these binding events), we decided to use t_2_ for our comparisons between sequences.

We then performed similar experiments with the same 8-nt gapped DNAs and mismatched DNA seals. Since the DNA length and the number of consecutive complementary base-pairs (bp) can determine the dissociation kinetics of a bound DNA, we expected that a seal DNA with a single-bp mismatch would bind for a shorter time than the fully complementary seal. We thus generated seals with a single internal mismatch by substituting cytosine for guanine (creating a G-G mismatch), and examined its effect on the dwell time; we also examined different positions for the mismatch.

We first examined the binding of an 8-nt seal DNA with a mismatch at position 3 (GS8-mis3G-T^ATTO647N,P28^; Fig. 2C, top); the use of this DNA yielded many transient binding events, which, as expected, were shorter than for the complementary DNA (Fig. 2C, middle). Dwell-time analysis confirmed this assessment and showed that a much shorter dwell time t_2_ (~2.2 s) when compared with the fully complementary seal (Fig. 2C, bottom); time t_1_ is also ~40% shorter, following the trend seen for t_2_.

We also examined a single G-G mismatch at position 5 (using DNA strand GS8-mis5G-T^ATTO647N,P28^, Fig. S2A); this mismatch was less disruptive to binding, yielding a dwell time t_2_ of ~ 9.5 s, which placed it between the fully complementary seal and the seal with a position-3 mismatch (Fig. S2B-C). Since both single-mismatch seals lead to a G-G mismatch and have the same number of G/C bases, the large difference in dwell times reflects the position of the mismatch within the seal, which has been shown to be important in single-molecule studies of short DNA duplexes (21).

### Increasing the sensitivity and throughput of sequence interrogation

To reduce the interrogation time for a single base, a fully complementary seal with a shorter dwell time is preferable, since this should lead to an overall faster sampling of binding. An approach to achieve this is to use a shorter seal; use of a shorter seal should also lead to a greater destabilising effect for a single mismatch, providing a clearer discrimination between the binding kinetics of a complementary seal and a single mismatch seal.

To test our ability to work with shorter seal DNAs, we reduced the seal size from 8 nt to 6 nt and compared a fully complementary seal (GS6-compl-T^ATTO647N,P28^) with a seal leading to single G-G mismatch at position 3 (GS6-mis3G-T^ATTO647N,P28^). Experiments with the fully complementary using 10-ms exposures resulted in time-traces showing >1,200 events of binding to the gapped-DNA (Fig. S3), with an average dwell time of t_2_ ~ 0.75 s (Fig. 2D). In sharp contrast, only 3 binding events were recorded for the single mismatch seal for the same volume of experiments, reflecting the fact that the mismatch seals bound on the gapped DNA for times much shorter than our exposure time.

To increase the seal binding rate, we also explored ways to increase the seal concentration; since the binding rate (on-rate) depended on the seal concentration, an increased concentration should increase the frequency of binding events. However, since the seals are fluorescently labelled, an increased seal concentration also increases the fluorescence background from the unbound probes, which prevents single-molecule detection at seal concentrations of >50 nM. We thus explored the reduction the background fluorescence of the unbound seals through contact-mediated quenching (i.e., static quenching) of the fluorophore (22–24); this strategy relies on the fact that the seals are short single-stranded DNAs that behave as dynamic random coils in solution, allowing the seal ends to come into close proximity. As the seal DNAs bind to their target to form a B-form dsDNA, the static quenching is removed, leading to bright fluorescence from the bound seal molecule.

To introduce static quenching in DNA seals, we used different quencher groups placed on the 3’ end of the seal, which is the most distal location from the primary fluorophore, which was placed on the 5’ end of the seal. First, we studied use of a seal with two ATTO647N fluorophores (one at the 5’ end and a second at the 3’ end), as it is known that two fluorophores can statically quenched each other when placed in close proximity (23); we indeed observed large fluorescence quenching for unbound seals, allowing the observation of single immobilised DNAs even at concentration of 500 nM of unbound 2xATTO647N seal (Fig. S4A). The presence of two terminal fluorophores did not hinder seal binding to the DNA; further, seal binding showed complete de-quenching upon binding to gapped-DNA (Fig. S5A-B; note that, in this experiment, the dequenching was observed via FRET from a nearby donor to the dequenched ATTO647N groups). Overall, the brighter fluorescence of the seal upon binding and the reduced background fluorescence due to contact quenching allowed for a ~10-fold increase in the allowable seal concentration (up to 300-500 nM).

Further, we tested the use of static quencher DABCYL (24) on the 3’ end of a seal carrying a ATTO647N at its 5’ end. In solution, unbound seals also showed large fluorescence quenching, although not as high in the 2xATTO647N case (Fig. S4B). Further, as with the 2xATTO647N strategy, seal binding showed substantial de-quenching upon binding to gapped-DNA (Fig. S5C-D).

The seal design with 2xATTO647N was chosen over the ATTO647N-DABCYL pair for further experiments, since the presence of two fluorophores on the seals lead to brighter fluorescence upon binding compared with a singly fluorophore labelled seals; the 2xATTO647N is also associated with slightly higher operational range of concentrations. The use of contact-mediated quenching should also be useful in other techniques relying on DNA-binding, such as DNA PAINT (15), FISH (25–27), and molecular beacons (22).

### Connecting single-protein binding to the sequence of a single DNA molecule

To develop ways to link a kinetic phenotype to DNA sequence, we examined the sequence-dependent binding of a bacterial transcription factor (catabolite activator protein, CAP) to surface-immobilised dsDNA, and connected the affinity of CAP-DNA binding to the DNA sequence at the single-molecule level. CAP is a global transcriptional activator that regulates transcription in *E. coli* mainly by accelerating the rate of gene promoter opening by RNA polymerase (RNAP). In the presence of cyclic AMP, CAP recognises the two-fold symmetric 22-bp consensus binding sequence 5’-AAATGTGATCTAGATCACATTT-3’ with high affinity (CAPcons DNA; Fig. 3A-B).

**Figure 3.**
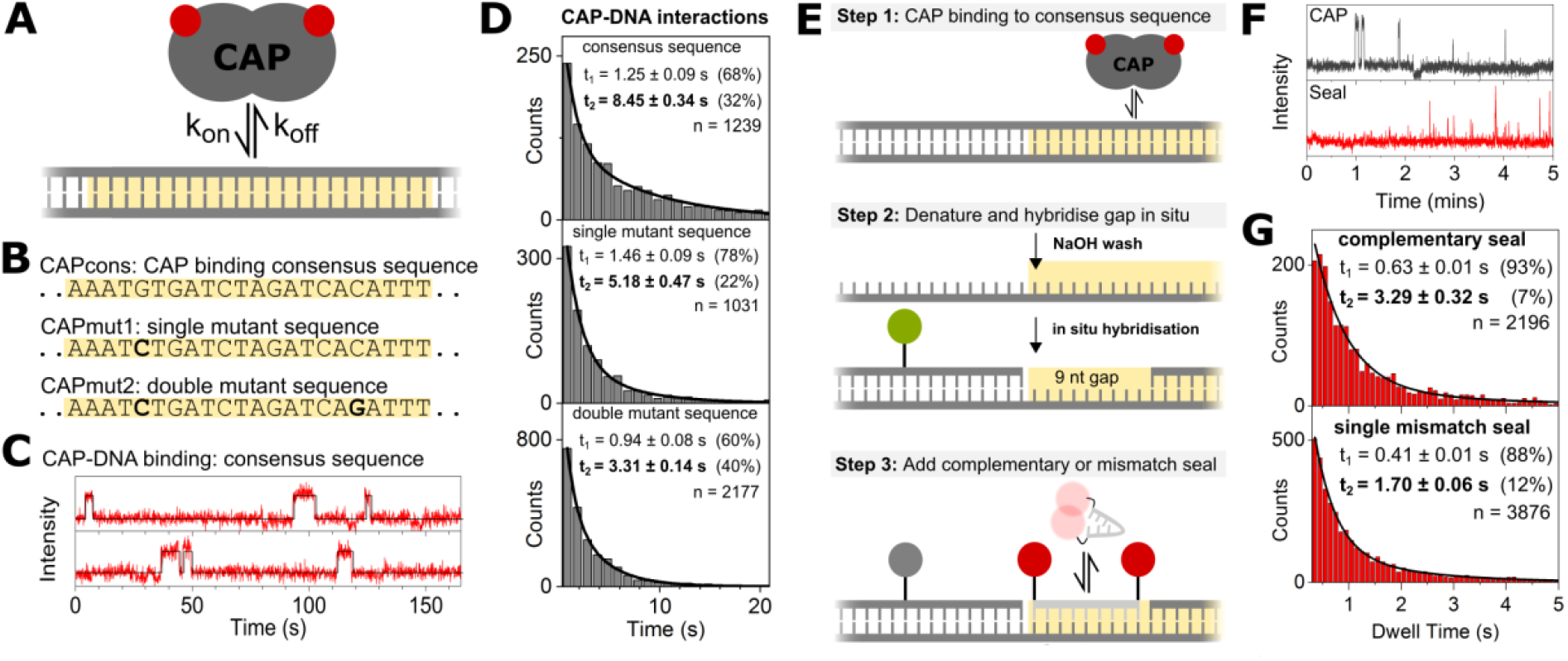
Linking the kinetics of protein-DNA interactions to DNA sequence at the single-molecule level: interaction of a transcriptional activator with its cognate DNA. A. Schematic of fluorescently labelled catabolite activator protein (CAP) binding to dsDNA with a 22-bp binding sequence for CAP (yellow shading). B. From top to bottom: symmetric 22-bp consensus CAP sequence binding sequence (CAPcons); single mutant CAP-binding sequence (CAPmut1); double-mutant CAP binding sequence (CAPmut2). C. Fluorescence intensity time-traces (red) for Alexa647-CAP binding to CAPcons DNA. HMM analysis identifies two different states in the traces: high intensity due to CAP binding, and low intensity due to absence of CAP. D. Dwell-time histogram for CAP binding to CAPcons (top), CAPmut1 (middle), and CAPmut2 (bottom) binding sequences. In all cases, the data fit best to double-exponential distributions that correspond to a long and short dwell time. E. Three-step assay for linking kinetic phenotype to sequence. <u>Step 1</u>: CAP binds to surface-immobilised CAPcons dsDNA. <u>Step 2</u>: dsDNA denatured by NaOH wash leaving only surface-immobilised bottom DNA strands, followed by gapped-DNA formation via in situ hybridisation of left-hand (fluorescently labelled) and right-hand flanking strands. <u>Step 3</u>: Fully complementary or single mismatch seal interrogates the gap sequence in the gapped-DNA. F. CAP binding trace (grey) in step 1 and complementary seal binding trace (red) in step 3 for a single DNA molecule on the surface. G. Dwell time histograms for seal binding in step 3 of the three-step assay. Top: Complementary seal. Bottom: Single mismatch seal (C-C mismatch at position 5).

To examine the effect of base substitutions on the kinetics of CAP binding to its consensus DNA sequence, we designed two CAPcos sequence variants (“mutants”; Fig. 3B). The first sequence (“single mutant” or “CAPmut1”) included a G/C pair conversion into a C/G pair at position 5 of the 22-bp sequence, a position known to make direct contacts with CAP. The second sequence (“double mutant” or “CAPmut2”) included the same change at position 5, as well as a C/G → G/C conversion at position 18, which is symmetric about the axis of symmetry in the CAPcons sequence.

To observe CAP binding to the CAPcons DNA site at the single-molecule level, we first immobilised a biotinylated Cy3B-labelled CAPcons DNA construct on neutravidin-coated PEGylated slides and localised the DNA using 532-nm excitation (see *Methods*). We then added 1.5 nM Alexa647-labelled CAP (see *Methods* for labelled CAP preparation), and observed CAP binding using 640-nm excitation in CAP-binding buffer (0.2 mM cAMP, 40 mM Tris-HCl pH 8, 100 mM KCl, 10 mM MgCl_2_, and 5% glycerol, plus an oxygen-scavenging system; see also *Methods*).

Under our buffer conditions, the binding of labelled CAP molecules to CAPcons DNA was transient, with CAP binding leading to a high level of fluorescence intensity, and subsequent CAP dissociation reducing the intensity to the background level (Fig. 3C); in fact, the traces resembled those observed for the transient binding of DNA seals to gapped DNA. Using a dwell-time analysis similar to that for transient DNA-DNA binding, we extracted dwell-times for CAP binding to its DNA site, and showed that the dwell-time distribution is described best by a bi-exponential fit (Fig. 3D, top); the longest lifetime t_2_ ~ 8.45 ± 0.34 s reflects the average dwell time for stable CAP binding to the full CAPcons DNA site, and corresponds to a CAP dissociation rate k_off_ of ~0.12 s^-1^. Further, analysis of the frequency of our binding events (Fig. S6) allowed us to determine that the CAP-DNA association rate constant k_on_ is ~9.73 x 10^6^ M^-1^s^-1^, which corresponds to a binding rate of 0.012 s^-1^ and a K_d_ of ~12 nM.

Use of the single-mutant CAPmut1 DNA site also led to similar transient events (Fig. S7A), but with a significantly shorter dwell time t_2_ ~ 5.2 s (Fig. 3D, middle); considering that k_on_ was 8.74 x 10^6^ M^-1^s^-1^, the single mutation resulted in a K_d_ ~ 22 nM, hence a ~2-fold loss in affinity. Further, CAP binding to the double-mutant CAPmut2 DNA site exhibited a further reduced dwell time (Fig. S7B), with a t_2_ ~ 3.3 s (Fig. 3D, bottom), a k_on_ of 10.24 x 10^6^ M^-1^s^-1^ and a K_d_ of ~ 30 nM. These results clearly establish that we can directly detect the changes in binding affinity and dissociation/association kinetics associated with one or more base changes in a protein-binding DNA sequence.

We then proceeded with linking the kinetic phenotype (here, the CAP-binding kinetics) associated with a *single* DNA molecule to the sequence of the *same* single DNA molecule; this linking required that we examine the same field of view of single DNA molecules on our microscope slide. We first conducted the single-molecule CAP-DNA binding experiments for step 1 of the assay (Fig. 3E, step 1), where CAP was observed to bind to CAPcons with t_2_ ~ 8.7 ± 0.3 s (Fig. S8), a value essentially identical to that obtained when we solely evaluated CAP-DNA binding (Fig. 3D, top).

For step 2 (Fig. 3E), the dsDNA was denatured and washed using 25 mM NaOH, removing the protein and the top DNA strand, and leaving only the biotinylated bottom DNA strand on the surface; subsequently, two DNA strands (complementary to sequences left and right of the sequence that will ultimately form the ssDNA gap) were hybridised in-situ to the bottom strand to form an immobilised gapped DNA featuring a 9-nt ssDNA gap, which was identical to the first 9 bases of the 22-bp CAPcons sequence.

In step 3, we performed transient DNA binding measurements for the same field of view used to image CAP binding in step 1 (Fig. 3E, step 3). First, we performed the entire assay using 100 nM fully complementary 9-nt seal DNA (GS9-CAP-T^AT647N,P21,P29^), recorded time-traces, and linked the CAP-DNA binding events to the DNA-DNA binding events for the same molecule (see example in Fig. 3F). The fully complementary seals yielded a dwell time of t_2_ = 3.29 ± 0.32 s (Fig. 3G, top). This time is shorter to those seen earlier in the paper (Figure 2), mainly reflecting the large difference in the G-C content of the gaps examined (22%, much lower than the 66% G-C content for 8-nt gapped DNA in Figure 2B-C), which reduces the free energy of binding by a ΔΔG of ~2.6 Kcal/mol (28).

We then repeated all 3 steps using 100 nM of a seal DNA with a single base-pair mismatch (GS9-CAP-mis5C-T^AT647N,P21,P29^, resulting in a C-C mismatch at position 5) for step 3, which led to a dwell time of t_2_ = 1.70 ± 0.06 s (Fig. 3G, bottom), almost half that of the fully complementary seal. Here, as previously shown, the fully complementary seal and the seal with a single mismatch can be distinguished on the basis of their dwell times of the immobilised CAPcons DNA.

### Mixed DNA sequences on a surface

All previous experiments were conducted using only a single DNA sequence immobilised on the surface; however, the implementation of Gap-seq will employ libraries of hundreds to thousands different DNA sequences, each represented by several DNA molecules placed randomly on the same glass coverslip. Notably, this experimental design assesses both complementary and mismatched sequences in the *same* field of view under the *same* experimental conditions (e.g., temperature), allowing any changes in kinetics to be directly linked to the sequence with minimal interference from external factors.

To test our ability to work with libraries of sequences, we mixed two solutions of molecules representing two different DNA sequences (namely, CAPcons and CAPmut1 sequences), and immobilised them randomly on the surface. To obtain the ground truth regarding the actual sequence of each individual DNA molecule, the CAPcons DNA was labelled (prior to immobilisation) with green fluorophore Cy3B, and the CAPmut1 DNA was labelled with red fluorophore Cy5 (Fig. 4A); this way, images of a field of view of a mixed DNA sample upon 532-nm and 640-nm excitation were sufficient to report the location of the molecules with different sequences (Fig. 4B).

**Figure 4.**
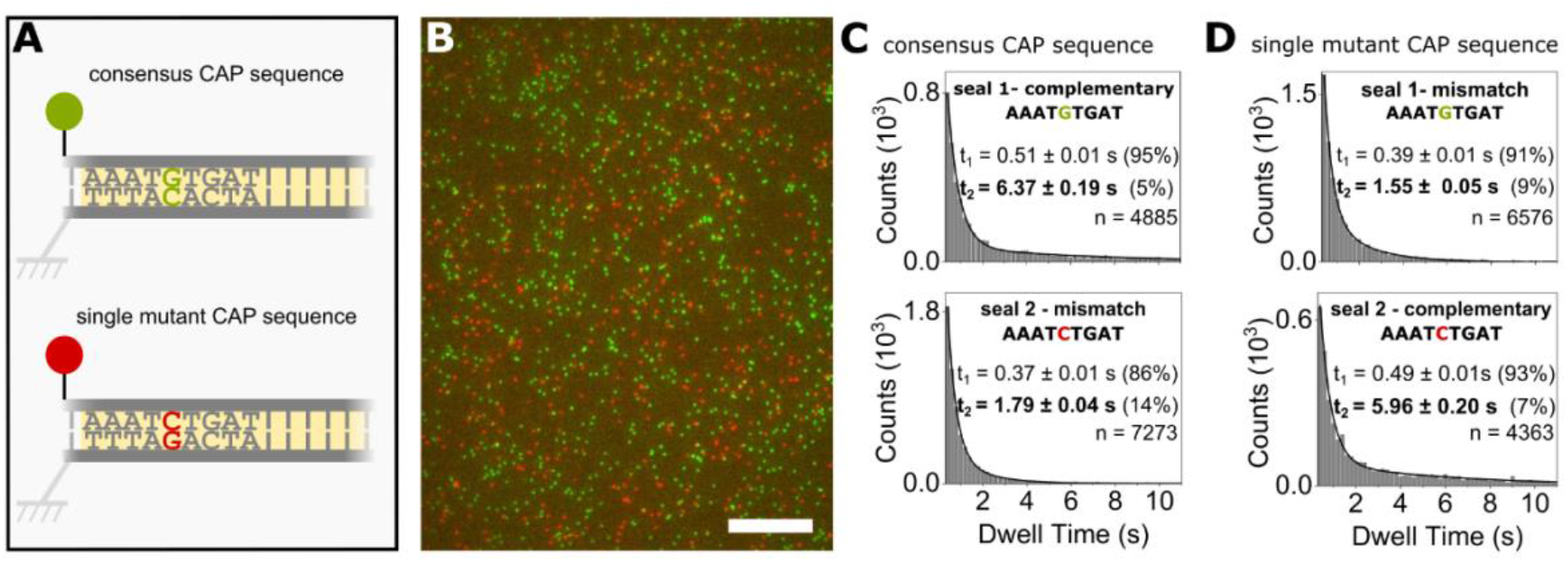
Distinguishing between molecules with different DNA sequences immobilised on the same surface. A. Top: CAPcons sequence labelled with Cy3B. Bottom: CAPmt1 binding sequence (G-C pair at position 5 flipped) labelled with Cy5. B. Microscope field of view showing dsDNA molecules featuring the CAPcons sequence (green) and the CAPmut1 sequence (red) randomly immobilised on the surface. Scale bar, 10 μm. C. Dwell-time histograms for seal binding to the CAPcons sequence. Top: Complementary seal. Bottom: Single mismatch seal (C-C mismatch at position 5). D. Dwell-time histograms for seal binding to the CAPmut1 sequence. Top: Single mismatch seal (G-G mismatch at position 5). Bottom: Complementary seal.

To demonstrate the ability to examine the fluorescence signatures (which act as a proxy for a kinetic phenotype) of mixed samples and connect them to their DNA sequence, we followed the same steps needed to link single-molecule phenotypes to DNA sequence (Fig. 3E), with the exception that Step 1 involved just the localisation of the DNAs (since CAP binding to the same DNA sequences had already been demonstrated; see Fig. 3D, top and middle). After gapped-DNA formation in step 2, we interrogated the immobilised DNAs in step 3 using a seal fully complementary to CAPcons (GS9-CAP-T^AT647N,P21,P29^, “seal 1”). The dwell time of seal 1 on CAPcons molecules was t_2_ ~ 6.4 s (Fig. 4C, top), whereas the dwell time of seal 1 on CAPmut1 molecules was much reduced, with a dwell time of t_2_ ~1.55 s (Fig. 4C, bottom). This 4-fold dwell-time reduction for seal 1 as we move from CAPcons to CAPmut1 provides strong evidence of the large effect of a single base-pair mismatch on DNA-hybridisation kinetics, and indicates how this approach can be used to sequence the ssDNA gap.

We then repeated our experiments by using in step 3 a different seal (seal 2; GS9-CAP-mis5C-T^AT647N,P21,P29^), which was fully complementary to CAPmut1; as a result, seal 2 bound to CAPmut1 much more stably that seal 1, and yielded a dwell time t_2_ ~ 6 s (Fig. 4D, bottom). On the other hand, seal 2 had a single base-pair mismatch (a G-G mismatch) to position 5 within the 9-nt gapped DNA of CAPcons, leading to a reduced dwell time of t_2_ ~ 1.8 s (Fig. 4D, top). As previously, a large reduction (~3.8-fold) in the dwell time can be detected for a single seal having just a single base-pair mismatch.

In conclusion, both experiments using mixed DNA sequences have shown that transient binding of short DNAs can detect single base changes in the sequence of surface-immobilised DNA, paving the way for future studies of protein-DNA interactions using libraries of different DNA sequences.

## DISCUSSION

Our work establishes the ability to extract DNA-sequence information via detection of transient binding of short DNA molecules to single surface-immobilised DNA molecules featuring short gaps, identifying the presence of a single base-pair mismatch for gaps of 6-9 nt. We also show how to link DNA-sequence information to protein-DNA interaction kinetics at the single-molecule level by looking at the effects of DNA substitutions on the DNA binding of a bacterial transcription factor.

Our work paves the way for an assay that links single-molecule kinetic phenotypes to short randomised sequences (5-6 nt) represented in moderate-size libraries (up to 4096 sequences). Such complexity will offer substantial insight on how DNA sequence impacts the structure, interactions, and reaction kinetics of many systems involving nucleic acids.

The connection of a single-molecule phenotype to a DNA sequence has also been encountered in studies of decoding of combinatorially modified nucleosomes (29), where histone modifications of nucleosomes were linked to DNA sequences in the genome, with the DNA being sequenced by sequencing-by-synthesis. In contrast, our work focuses on obtaining kinetic phenotypes on DNA, with the DNA sequence interrogation relying on transient single-molecule hybridization.

Our results provide insight on the kinetics of DNA hybridization, especially as it applies to short DNA duplexes (6-10 nt), similar to the ones used in DNA-PAINT (15), in biosensing assays for diagnostics (30), and in computational simulations that dissect DNA hybridization mechanisms (31). Our work also identified DNA modifications that suppress signals by unbound fluorescent species (a major limitation in single-molecule fluorescence applications (32)) without using sophisticated devices such as zero-mode waveguides (12).

### Design considerations

Currently, the optimum length of the DNA gap to be sealed by transiently bound DNA is 6-9 nt. A seal longer than 9 nt will yield dwells longer than 10 s, decreasing the number of binding and dissociation events, and increasing statistical noise; the effect of a single mismatch on the interaction kinetics will also be smaller. On the other hand, a seal shorter than 6 nt will be hard to capture in its gap-bound state (given the typical temporal resolution of single-molecule fluorescence instrumentation), and will limit the size of the library to 4^5^ sequences or less. It may, however, be possible to use a 5-mer seal (expected dwell time of 50-100 ms) using higher excitation powers and exposure times of 5-10 ms; notably, at that length, the discrimination between a complementary and a single-mismatch seal will be significantly enhanced. Use of 5-mer seals will also benefit from using modified nucleotides which can increase affinity for the complementary target (e.g., locked nucleic acids [LNA], modified bases or spermine monomers), and might also improve mismatch discrimination.

Since the effects of mismatches are most evident in internal positions of the DNAs corresponding to the gap (21), it will be best to insert the libraries at the middle 5 positions of a 9-mer; our results here clearly establish the ability to observe mismatches at such positions.

### DNA hybridization kinetics on gapped DNA substrates

Our results show that the dissociation of short DNAs from a DNA gap follows a double-exponential decay, pointing to the formation of two hybridized species with markedly different stability. Notably, studies of hybridization kinetics on non-gapped DNA (15, 21, 33, 34) have not observed such a double-exponential behaviour. We attribute our long dwell times to a fully hybridized DNA (stabilised by stacking both at the 5’ and 3’ ends, and by complementary binding to all of the template bases), and our short dwell times to partially hybridized DNA (that lacks stacking on either one or both of the ends). Notably, the shorter dwells times correspond better to the lifetimes observed previously in the absence of stacking, and may also correspond to frayed states of the strands flanking the gap. Such partially hybridized species have also been observed in computational studies using Ox-DNA (31), albeit these simulations involved hybridization of single-stranded DNA in the absence of a gap and its associated stacking-based stabilisation.

### Assay improvements

Further development of the assay will involve integration of microfluidics, sequential base reading, and faster data acquisition and analysis. Integration of automated microfluidics will eliminate the tedium associated with manual washing and sample-loading steps, and permit analysis of DNA libraries, which will require many fluid-delivery events to surface-immobilised samples.

Future work will use the dwell-time information (and, more generally, the single-molecule trace), to call the identity of a template base on a single molecule relying on the identification of the complementary binding behaviour vs. the mismatch binding behaviour for a single template base; this step will test sequentially all four bases per template position. These efforts can also leverage the binding rate (k_on_) of the seals, since it can help base identification (21). To read longer sequences (e.g., a 5-nt sequence) on the gapped DNA, we will examine the identity of each unknown bases sequentially, utilising 4 seal DNAs per base, with each seal DNA containing one of the 4 natural bases (to examine complementarity with the interrogated base), as well as either “universal bases” (such as nitroindole (35)) or “degenerate bases” (formed when a mixture of the four natural bases is used in one addition cycle during automated DNA synthesis), arranged to oppose some or all of the remaining nucleotide positions of the gap section.

There is also scope to reduce the acquisition time for a single base interrogation, which will in turn minimise issues related to thermal drift, instrument stability, and data storage. Faster acquisition will be helped by working with higher densities of gapped molecules, which in turn can be achieved by resolving closely spaced molecules via high-precision localisation (15). The data analysis will also benefit from automated time-trace selection and methods that determine the degree of binding without dwell analysis.

### Biomolecular systems and interactions amenable to Gap-seq

The Gap-seq approach will allow detail study of the sequence-dependence of processes involving proteins that read and process DNA in multiple steps (RNA polymerases, gene-editing enzymes), as well as proteins that bind and deform DNA (transcription factors, proteins that detect DNA damage, nucleoid-associated proteins); we are particularly interested in exploring how promoter sequences shape the complex kinetic landscape of transcription initiation (6), initial transcription (36), and promoter escape (37), all of which are central to the regulation of gene expression in all organisms. Extension to RNA-binding proteins is also possible, starting with monitoring a protein-RNA interaction, followed by reading the RNA sequence using RNA-DNA hybridization. Since the nucleic acid library can comprise RNA or modified DNA constructs that fold into aptamers (38), our approach offers a way to determine the sequence-specificity for protein-aptamer binding, and to generate improved aptamers for protein or biomolecular capture, labelling, and detection, as well as for drug discovery. Our methodology could also be used to study triplexes, i.e., the binding of a third strand to an immobilised duplex.

## ACKNOWLEDGEMENTS

We thank Dr Christof Hepp for help with the use of fiducial markers for drift correction, Mirjam Kümmerlin for helpful comments on the manuscript, and the MICRON Advanced Bioimaging Facility (supported by Wellcome Strategic Awards 091911/B/10/Z and 107457/Z/15/Z) for access to a Nanoimager microscope in their facilities. This work was supported by the Wellcome Trust (110164/Z/15/Z to A.N.K.), a UK EPSRC studentship (project 1733991, to R.A), and BB/R008655/1 (New and versatile chemical approaches for the synthesis of mRNA and tRNA) to A.H.E.S., T.B. and A.S.

## CONFLICT OF INTEREST STATEMENT

The work was performed using miniaturised commercial microscopes from Oxford Nanoimaging, a company in which A.N.K. is a co-founder and shareholder. R.A., H.S., A.H.E.S., A.S., T.B., and A.N.K. have financial interests in a patent application related to the work described in this manuscript.

## DATA AVAILABILITY

Single-molecule movies and fluorescence intensity time-traces will be available upon request.

