## Supplemental Figures for "Transient DNA binding to gapped DNA substrates links DNA sequence to the single-molecule kinetics of protein-DNA interactions"

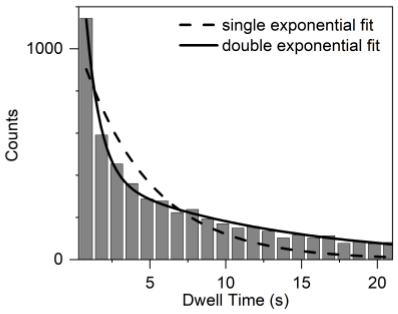


**Figure S1. Example of single- vs double-exponential decay fit for DNA seal bound times.** Dwell-time histogram for the fully complementary 8-mer binding to 8-nt gapped DNA fitted poorly to a single-exponential decay, while fitting much better to a double-exponential decay.

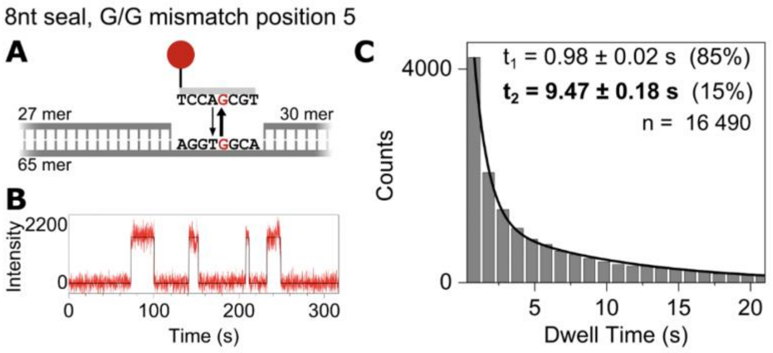


**Figure S2. Detecting a single base-pair mismatch via transient DNA binding: results for an 8-mer with a position 5 mismatch.**

A. Schematic of an 8-nucleotide (nt) seal forming a G-G mismatch at position 5 while binding to the 8-nt gapped-DNA.

B. Fluorescence intensity time-traces (red) for ATTO647N seal binding to gapped DNA. Movies recorded using 100-ms exposures. Traces are fitted (black) by Hidden Markov Modelling (HMM), identifying bound states and measuring the corresponding dwell times.

C. Dwell-time distribution for the bound states. The data fit best to double-exponential distributions that correspond to a long and short dwell time. n: number of binding events.

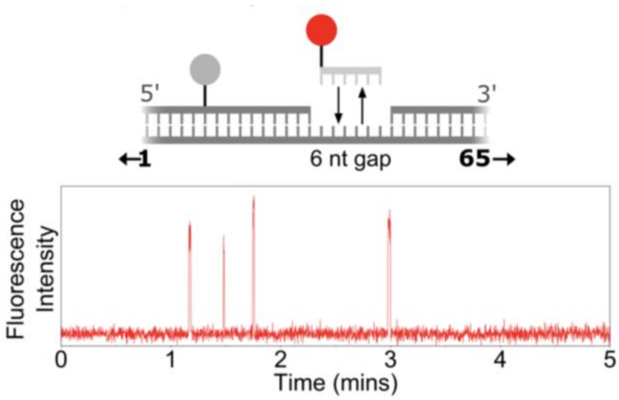


**Figure S3. Time-traces for 6-mer complementary seal.** GS6-compl-T$^{ATTO647N,P28}$, used at 30 nM, hybridises transiently to a 6-nt gapped-DNA. Movies were recorded using 10-ms exposures.

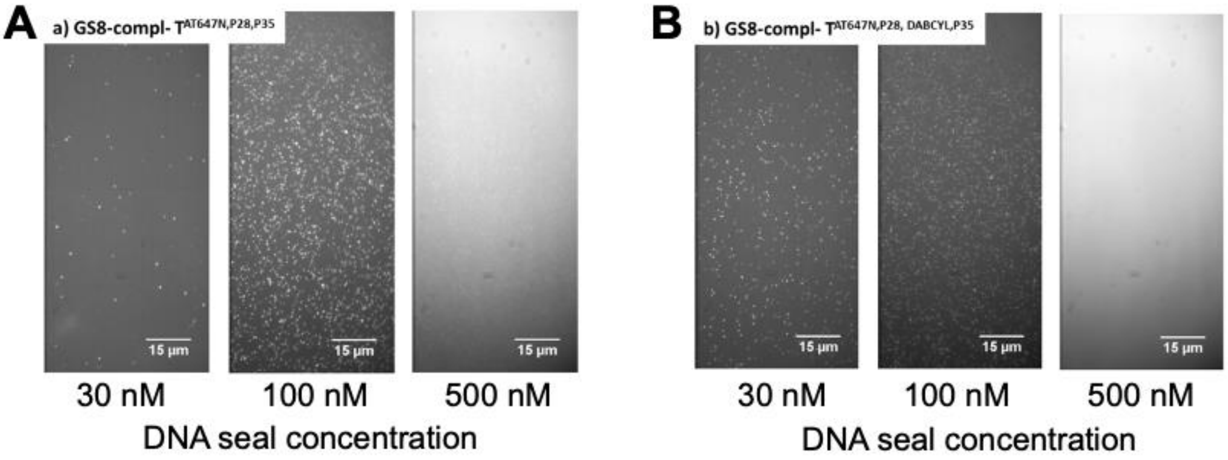


**Figure S4**: Microscope images (red emission channel) of seals and surface-immobilised 8-nt gapped-DNA and illuminated using 638-nm excitation.

A. GSeq-8compl-T$^{ATTO647N\ P28,P35}$ seal DNA added at 30 nM, 100 nM and 500 nM.

B. GSeq-8compl- T$^{ATTO647N\ P28,DABCYL,P35}$ seal DNA added at 30 nM, 100 nM and 500 nM.

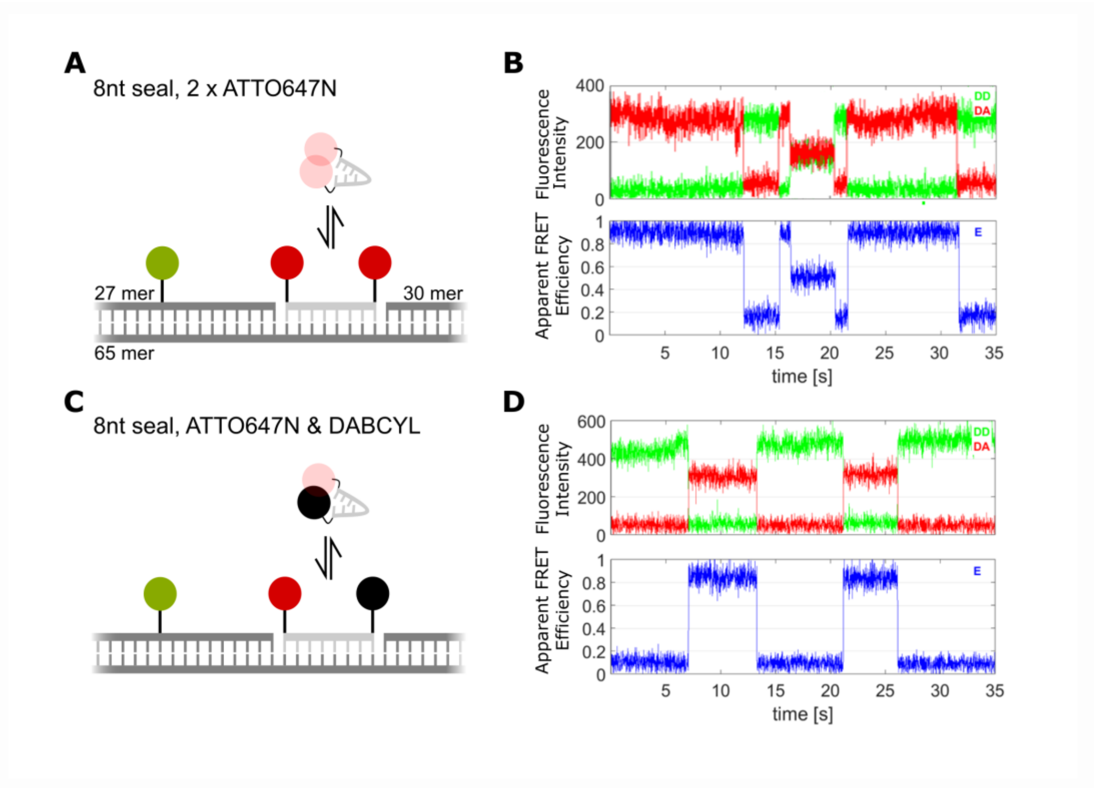


**Figure S5. Use of seal DNAs with engineered contact-mediated quenching of fluorescence**.

A. Schematic of the fluorescence quenching of GSeq-8compl-$T^{ATTO647N\ P28,P35}$ and dequenching upon binding to the gapped DNA. The dequenching will allow the two ATTO647N fluorophores to act as very efficient FRET acceptors (with apparent energy transfer efficiency of E*>90%) to the green donor fluorophore (Cy3B, located 10 bp [i.e., ~3.5 nm] away from the proximal ATTO647N probe) on the left flanking DNA of the gapped DNA.

B. Fluorescence intensity and FRET time-traces of 100 nM GSeq-8compl-$T^{ATTO647N\ P28,P35}$ seal transiently binding to an 8-nt gapped-DNA construct. DD, donor excitation upon donor emission; DA, donor excitation upon acceptor emission. As expected, GSeq-8compl-$T^{ATTO647N\ P28,P35\ FRET}$ binding leads to high apparent FRET efficiencies (E* >90%); bleaching of the donor-proximal fluorophore (at ~17 s) reduces E* to 50%, corresponding to FRET from the donor to the distal acceptor (17 bp [i.e., 6-7 nm] away). Exposure time, 100 ms.

C. Schematic of the fluorescence quenching of GSeq-8compl-$T^{ATTO647N\ P28,\ DABCYL\ P35}$ and dequenching upon binding to the gapped DNA. As in panel A, the dequenching allows the ATTO647N fluorophore to act as a very efficient FRET acceptor to the green donor fluorophore, leading to E*>90%.

D. Fluorescence intensity and FRET time-traces of 100 nM GSeq-8compl-$T^{ATTO647N\ P28,\ DABCYL\ P35}$ seals transiently binding to an 8-nt gapped-DNA construct. Binding leads to high apparent FRET efficiencies (E* of >90%). Exposure time, 100 ms.

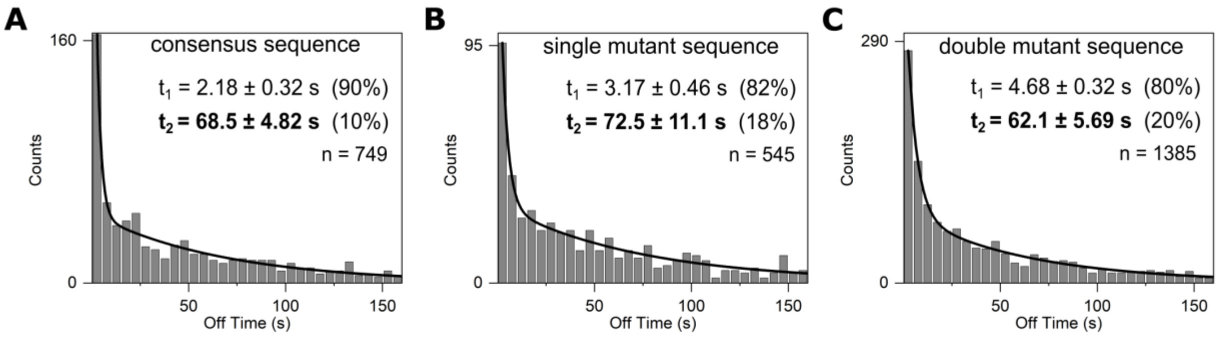


**Figure S6. CAP-DNA interactions: dwell-time distribution of unbound states (to get on-rate)**

Off-time distributions for CAP-DNA interactions with the consensus sequence (panel A), single mutant sequence CAPmut1 (panel B) and the double mutant sequence CAPmut2 (panel C). In all cases, the data fit best to double-exponential distributions that correspond to a long and short dwell time. n: number of binding events.

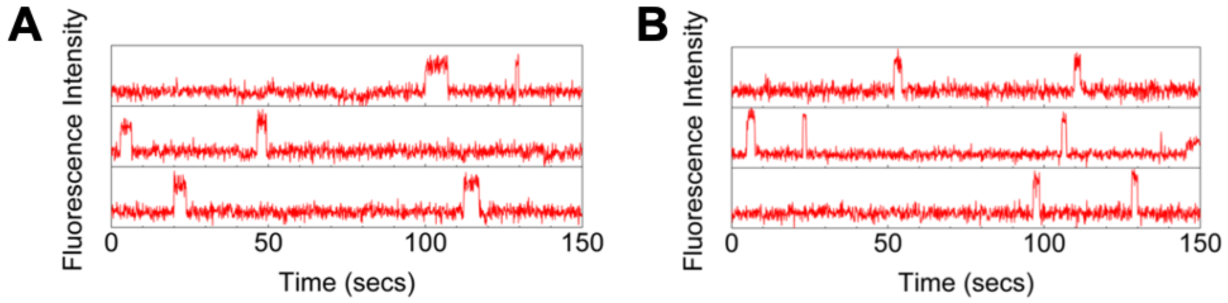


**Figure S7. Time-traces of CAP binding to mutant sequences.** Description as in Fig. 3C.

A. Three representative time-traces showing binding of CAP to the CAPmut1 sequence.

B. Three representative time-traces showing binding of CAP to the CAPmut2 sequence.

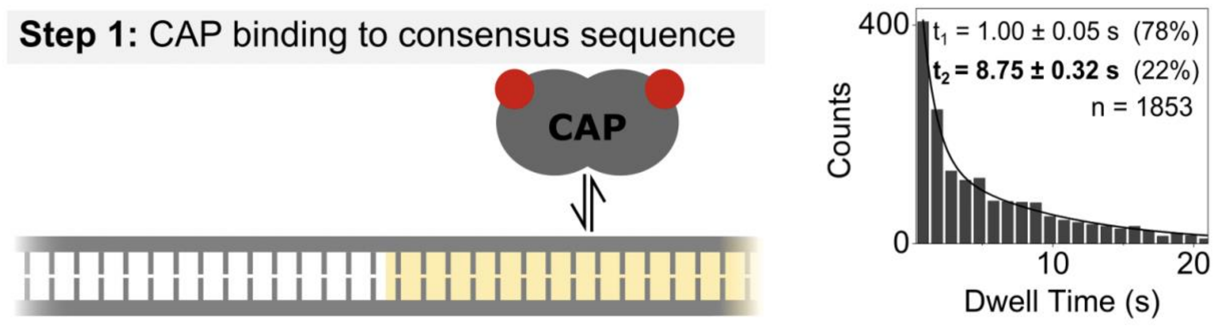


**Figure S8. CAP-binding histogram in step 1 of the full three-step assay.** Left, schematic of the CAP-DNA interaction. Right, dwell-time histograms for CAP bound to the CAPcons DNA site. The display of the exponential analysis is as in Fig. 3D.
